# Krumholzibacteriota and Deltaproteobacteria contain rare genetic potential to liberate carbon from monoaromatic compounds in subsurface coal seams

**DOI:** 10.1101/2023.07.10.548433

**Authors:** Bronwyn C. Campbell, Paul Greenfield, Se Gong, David J. Midgley, Ian T. Paulsen, Simon C. George

**Affiliations:** Environment Business Unit, Commonwealth Scientific and Industrial Research Organisation (CSIRO), Floreat, WA, 6014, Australia; School of Natural Sciences, Macquarie University, North Ryde, NSW, 2109, Australia; Energy Business Unit, Commonwealth Scientific and Industrial Research Organisation (CSIRO), Lindfield, NSW, 2070, Australia

## Abstract

Biogenic methane in subsurface coal seam environments is produced by diverse consortia of microbes. Although this methane is useful for global energy security, it remains unclear which microbes can liberate carbon from the coal. Most of this carbon is relatively resistant to biodegradation, as it is contained within aromatic rings. Thus, to explore for coal-degrading taxa in the subsurface, this study used coal seam metagenomes to reconstruct important metagenome-assembled genomes (MAGs) using a key genomic marker for the anaerobic degradation of monoaromatic compounds as a guide: the benzoyl-CoA reductase gene (*bcrABCD*). Three taxa were identified with this genetic potential. The first was a novel taxon from the Krumholzibacteriota phylum, which this study is the first to describe. This Krumholzibacteriota sp. contained a full set of genes for benzoyl-CoA dearomatisation, in addition to other genes for anaerobic catabolism of monoaromatics. Analysis of Krumholzibacteriota MAGs from other environments revealed that this genetic potential may be common within this phylum, and thus they may be important organisms for the liberation of recalcitrant carbon on a global scale. Further, two taxa from the Deltaproteobacteria class were also implicated in monoaromatic degradation; two geographically unrelated *Syntrophorhabdus aromaticivorans* MAGs, and a Syntrophaceae sp. MAG. Each of these three taxa are potential rate-limiting organisms for subsurface coal-to-methane biodegradation. Their description here provides an understanding of their function within the coal seam microbiome, and will help inform future efforts in coal bed methane stimulation, anoxic bioremediation of organic pollutants, and assessments of anoxic carbon cycling and emissions.

**Importance:** Subsurface coal seams are highly anoxic and oligotrophic environments, where the main source of carbon is “locked away” within aromatic rings. Despite these challenges, biogenic methane accumulates within many of these coal seams, which implies that the coal seam microbiome can “unlock” this carbon source *in situ*. For over two decades, researchers have been working to understand which organisms are responsible for these processes. This study provides the first descriptions of these organisms. Here, we report metagenomic insights into the liberation of carbon from aromatic molecules typically found within coal, the degradation pathways involved, and descriptions of the Krumholzibacteriota sp., *Syntrophorhabdus aromaticivorans*, and Syntrophaceae sp. that contain this genetic potential. Additionally, this is the first time that the Krumholzibacteriota phylum has been implicated in anaerobic dearomatisation of aromatic hydrocarbons. This potential is identified here in numerous taxa within the phylum from other subsurface environments, implicating Krumholzibacteriota in global-scale carbon-cycling processes.

## INTRODUCTION

The global energy transition from hydrocarbons to renewables requires low-emissions fuels for facilitating energy security during this shift. For this transition fuel need, methane is one such lower-emission alternative to coal (1). Methane gas provides a dispatchable source of energy that, unlike coal, produces neither particulates nor harmful nitrous and sulfur oxides during combustion (2, 3). Thus, significant research interest exists in enhancing rates of methane gas production in subsurface coal by using the coal seam microbiome (4–6).

Within the coal itself, carbon is primarily (∼60 to 100 %) contained within aromatic rings, which increase in abundance with thermal maturity (7). Thus, taxa capable of aromatic ring degradation may be the dominant contributors of carbon to the coal seam microbiome, especially in more thermally mature coals. Over the last decade, a good understanding has been formed of the range of microbes that occur in subsurface coal seams, which are typically characterised by a few relatively abundant taxa and a modest tail of rarer taxa (4, 8, 9). Coal seam microbiome studies have also revealed that the dominant organisms, such as taxa within *Desulfuromonas* and *Desulfovibrio*, are not involved in aromatic degradation but rather degrade simpler intermediates (8, 10). Aromatic-degrading taxa therefore probably occur within the rarer taxa.

Identification of aromatic-degrading taxa is important both for enhancing applied outcomes such as industrial gas production, and for understanding carbon mobilisation from the fossilised geosphere. Indeed, the research effort to identify these taxa has been ongoing since the early papers in the subsurface coal microbiology field (11). Some taxa have previously been identified with the genetic tools for aerobic aromatic ring degradation within coal seams (12–14), however, aerobic pathway contributions are likely restricted to very shallow regions of meteoric water infiltration, since subsurface coal seams are overall highly anoxic environments (14). Within other anoxic hydrocarbon-degrading environments, such as oil reservoirs (15), the catabolism of aromatic substrates progresses at least in part via intermediate aromatic compounds. Although a wide range of monoaromatic biodegradation pathways are known, these pathways are dependent upon a relatively small range of central intermediate aromatic compounds. One intermediate aromatic compound in particular, known as benzoyl-CoA, is central to the anaerobic degradation of a particularly wide range of monoaromatics (16). Further catabolism from this central intermediate requires the benzoyl-CoA reductase enzyme, which is responsible for the first dearomatisation steps of the ring (17). This enzyme for ring cleavage is a crucial step for accessing the carbon contained within the ring structures, as the thermodynamic stability of the aromatic ring structure renders it highly resistant to degradation (15). The ability to encode this enzyme and those for the proceeding metabolites in the benzoyl-CoA pathway (depicted in Figure 1) would provide organisms with access to much of the carbon locked up in the organic substrates present in coal.

**Figure 1.**
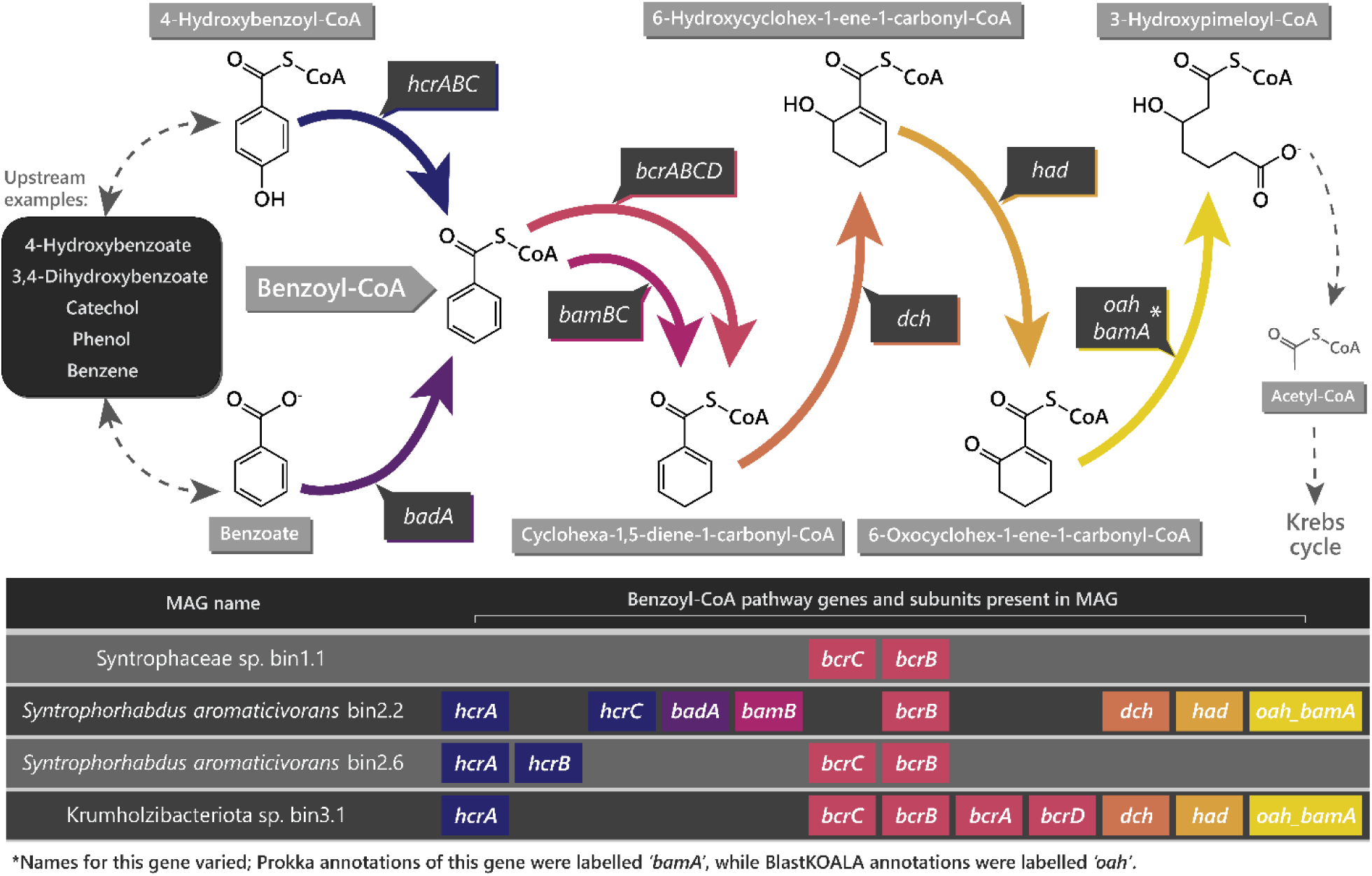
Benzoyl-CoA (Coenzyme A) pathway genes and subunits, from 4-hydroxybenzoyl-CoA and benzoate to 3-hydroxypimeloyl-CoA, and their presence in the selected contig bins (MAGs). Benzoyl-CoA reductase genes for two enzyme systems are displayed: the class-I, ATP-dependant *bcrABCD* (associated with facultative anaerobes), as well as the class-II, ATP-independent *bamB* (associated with obligate anaerobes) (15, 48). See Supplementary Table S2 for number of occurrences of each gene in the MAGs and metagenomes. The displayed benzoyl-CoA pathway was adapted from www.genome.jp/kegg/.

To date, numerous studies have attempted to identify aromatic-degrading taxa in coal seams using a range of strategies. Some of these have included the enrichment of microbes capable of degrading aromatic compounds, using either a variety of mono- and polyaromatic compounds or organic matter of differing maturities as sole sources of carbon (18, 19). These studies identified putative aromatic-degraders among the enriched taxa, but also obscure primary aromatic-degraders amid the numerous taxa subsequently enriched on their downstream degradative products.

Metagenome-assembled genomes (MAGs) have also been used to identify aromatic-degrading taxa by using known genes required for hydrocarbon degradation, performed alongside methods such as bio-orthogonal non-canonical amino acid tagging (BONCAT) to differentiate active from inactive cells (12, 13). Within the active taxa, this strategy has identified a range of genes used for rearranging the substituents of the aromatic ring in peripheral pathways above the benzoyl-CoA intermediate, for example genes for catabolism of ethylbenzene to acetophenone in Chlorobi and *Geobacter* taxa (12). Although these studies identify an abundance of genes and taxa actively involved in the coal-to-methane degradation pathways, the specific anaerobic monoaromatic- degrading genes identified are not involved in catalysing dearomatisation reactions or capable of liberating carbon from their targeted substrates (15, 20, 21). Consequently, the identification of taxa containing the genes for dearomatisation of these monoaromatic substrates, such as the benzoyl-CoA reductase gene, remains an unanswered and critical step for understanding the *in situ* liberation of carbon from coal.

Recently, Syntrophorhabdaceae and Syntrophaceae were implicated as potentially important coal-degrading families using the linear discriminant analysis effect size statistical method on a group of algal-amended coal seam microbiomes from the Powder River Basin, USA (22). Syntrophorhabdaceae is a monotypic family, of which *Syntrophorhabdus aromaticivorans* is presently the sole described species (23). *S. aromaticivorans* has previously been identified within an enriched Surat Basin (Australia) coal seam microbial community, which had likely responded to the increased surface area of the provided coal that had undergone solvent extraction (24). Outside the coal seam environment, a *S. aromaticivorans* isolated from an anaerobic digester was demonstrated to anaerobically catabolise multiple monoaromatic substrates to acetate, to utilise a model organic electron acceptor, and was proposed to be capable of interspecies electron transfer in partnership with a hydrogenotrophic methanogen (23). *S. aromaticivorans* is thus a promising candidate for monoaromatic degradation capabilities within the coal seam environment. In contrast to this, Syntrophaceae spp. are more commonly associated with aliphatic degradation than aromatic degradation, however, some taxa within the family can utilise a limited range of more labile monoaromatic substrates (25, 26). Although Syntrophaceae spp. and *S. aromaticivorans* have been implicated as taxa that may be important for *in situ* coal degradation, there has been no demonstration that strains from this environment have the genetic tools required to access aromatic compounds or culture-based demonstrations of their activity against aromatic constituents of coal.

One alternative strategy to those outlined above is to ‘mine’ metagenomic data for specific aromatic-degrading genes, and then use binning techniques to reassemble genomes of potential aromatic-degraders. Accordingly, this study aimed to identify aromatic-degrading taxa by mining metagenomic data from Australia and North America through first identifying assembled DNA sequences that contained benzoyl-CoA reductase gene subunits, and then using these as references to assemble metagenome-assembled genomes (MAGs) for the identification of these taxa and their capabilities.

## MATERIALS AND METHODS

### Sources of metagenomic DNA

For further details of metagenomic sequence data sourcing, see Campbell et al. 2022 (27). Briefly, whole-genome shotgun sequences selected for use were required to: (1) be from a subsurface coal seam, (2) state the coal seam or associated geological basin, (3) have been sequenced using Illumina, and (4) be available as unassembled sequence reads. Four metagenomes from Australia and nine from the USA were found to be suitable (Supplementary Table S1).

### Reassembly, annotation and detection of benzoyl-CoA reductase gene subunits

All metagenomes were downloaded as unassembled reads. These reads were error-corrected using Blue v2 (28) prior to assembly using SPAdes v3.13.2 with the meta flag (29). Resultant contigs were annotated using prokkaMeta v1.14.5 (30), and these annotations were then used to identify the various subunits of benzoyl-CoA reductase within the contigs.

### Selection of contigs with benzoyl-CoA reductase gene subunits

Contigs with benzoyl-CoA reductase gene subunits were interrogated using the Prokka annotations for other genes in the benzoyl-CoA pathway. Contigs selected for further analysis contained at least one benzoyl-CoA reductase gene subunit (*bcrABCD*), were of lengths greater than seven kilo–base-pairs, and had coverages greater than five.

### Generation of trimer signatures, correlations, bin quality control, and accessioning

In order to identify other contigs within the metagenome from the same taxon, a trimer approach was used (31). In brief, a custom Python script was used to count the proportion of each of the 64 possible trimers in all contigs containing at least 1000 base pairs. The trimer signature for those contigs containing benzoyl-CoA reductase gene subunits was then used as a reference, and the trimer signatures for all contigs in the metagenome were examined against these reference contigs using a Pearson correlation coefficient in SciPy 1.7.3 (scipy.org). Those contigs with Pearson’s R values greater than 0.95 were collected into bins for quality control. As a subsequent step in bin quality control, the coverages of each contig within the bin were inspected manually and contigs with aberrant coverage were removed. Each bin was then subject to inspection for completeness and contamination using CheckM v1.1.3 (32).

### Characterisation of the genomic content of the bins

Summary contig statistics (total bin length, total contigs, mean contig length, median contig length, N50, maximum contig length, and GC-content) were determined using a custom Python script. The bins were then submitted to online tools to further characterise the genomic content of each bin. For use with KEGG Mapper (genome.jp/kegg/mapper), the Prokka-annotated amino acid files were submitted to the BlastKOALA v2.2 online tool and run against the “species_prokaryotes” database (33). The Prokka-annotated amino acid files were also used to determine which transport proteins were present in each bin, by submission to the TransportDB 2.0 TransAAP (membranetransport.org) for transporter annotation (34). In order to characterise the carbohydrate active enzymes, dbCAN HMMdb v10.0 (bcb.unl.edu/dbCAN2) was run on the unannotated contigs within each bin (35). Similarly, the CRISPR sequences and *cas* genes were identified in the contigs within each bin using CRISPRCasFinder online (crisprcas.i2bc.paris-saclay.fr; (36)). Resultant data from these tools were summarised and used to characterise the genomes in each bin and estimate their ecological roles within the coal seam environment.

### Identification of the bins

As all bins lacked 16S ribosomal DNA (rDNA) regions, the elongation factor G genes were used to identify the closest known relatives of each bin. For this, BLASTX searches of the non-redundant protein database were used (https://blast.ncbi.nlm.nih.gov/Blast.cgi). Subsequently, putative identities from elongation factor G were compared with BLASTN searches of 16S rDNA sequence data obtained from each metagenome using the Earth Microbiome Project V4 primers with Kelpie (37–39). This manual comparison between the elongation factor G taxonomy and Kelpie-derived 16S rDNA data was used to identify putative taxa at the level of family or lower, with a minimum per-taxon abundance of 10 applied for the 16S rDNA data. In addition, all 16S rDNA sequences generated using Kelpie were then compared to the Coal Seam Microbiome (CSMB) reference set, using USEARCH v11.0.667 at 97 % identity, to determine any associated CSMB reference taxa (8, 40).

Where close elongation factor G sequence % identities were not found with BLASTX, the phylogeny of the bin was estimated against other Prokka-annotated representative genomes using multilocus sequence analysis (MLSA). The common housekeeping genes selected for this were restricted to those present within both the bin and the majority of the representative genomes. The constructed fasta files of the selected housekeeping genes were aligned and clustered in Mega v11.0.13 (41) with the ClustalW function, and edited as Newick tree format files in FigTree (http://tree.bio.ed.ac.uk/software/figtree/).

### Figures

Final editing of each figure was performed using Inkscape v1.1.2 or Adobe Illustrator v26.0.2.

### Data availability

Final contig bins, Prokka nucleotide annotations, and Kelpie-generated 16S rDNA sequences were uploaded to the CSIRO Data Access Portal (https://data.csiro.au/collection/csiro:54006). An overview of the methods can be found in Supplementary Figure S1.

## RESULTS

### Distribution of benzoyl-CoA reductase genes across the examined metagenomes

Subunits of the benzoyl-CoA reductase gene (*bcrABCD)* were detected in the Prokka annotations of all examined metagenomes except for Central Appalachian 89 (Supplementary Table S2). The greatest number of benzoyl-CoA reductase subunits were detected in the Powder River 40 metagenome, with 43 subunits observed. In contrast, aside from the Central Appalachian 89 metagenome, the Bowen 3 metagenome contained the fewest number of benzoyl-CoA reductase subunits (4 subunits; Supplementary Table S2). Overall, subunits *bcrB* and *bcrC* were identified far more often than subunits *bcrA* and *bcrD*; no *bcrA* or *bcrD* subunits were detected in the Australian metagenomes, and in the Powder River Basin metagenomes *bcrBC* were an order of magnitude more abundant than *bcrAD* (215 total compared to 23 total, respectively).

Twenty-one bins were obtained that contained benzoyl-CoA reductase gene subunits, and comparison of these indicated three groupings of similar genomes (Supplementary Table S3). From these groupings, four bins were characterised in detail (bins 1.1, 2.2, 2.6 and 3.1; Figure 1; Table 1). The remaining 17 bins were determined to be chimeric, as they were too large to represent a single genome, or to have unacceptably low CheckM contamination scores (Supplementary Table S3).

**Table 1:**
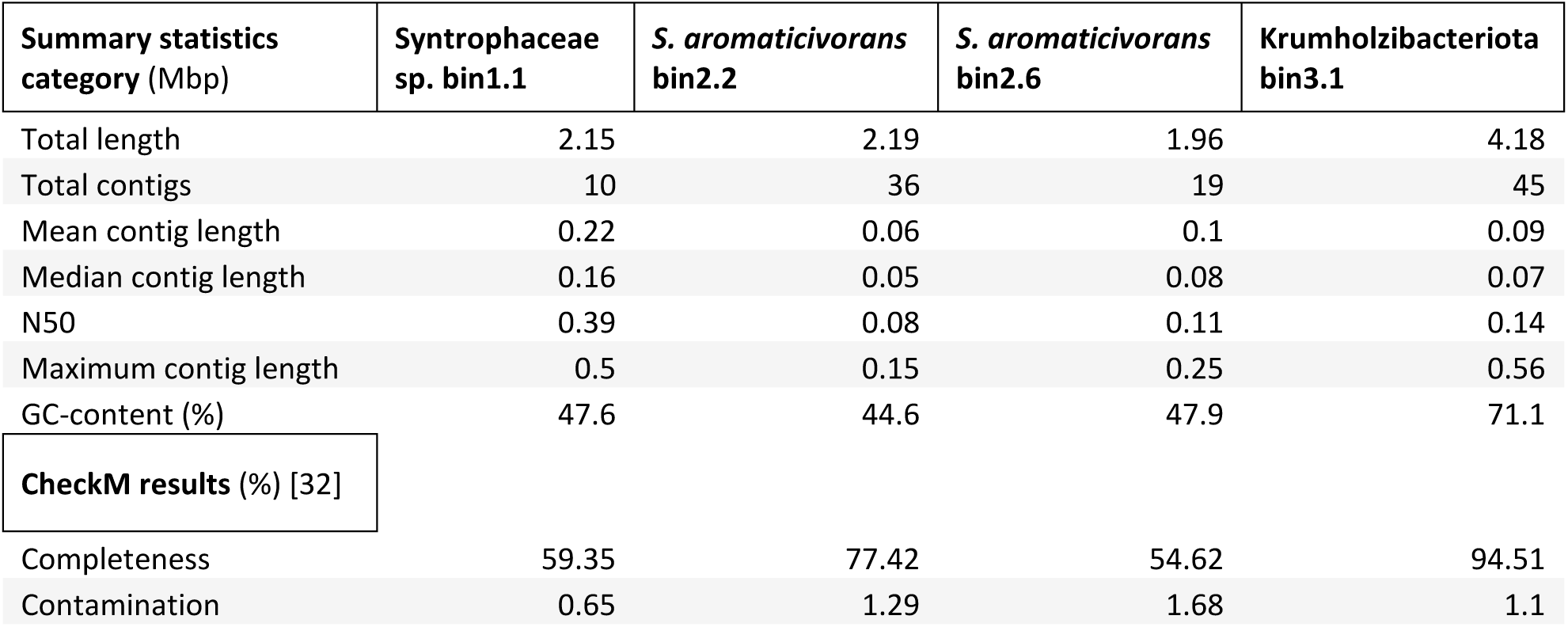
B**i**n **statistics**.

### Phylogeny of the four contig bins that were chosen for further analysis

16S rDNA sequences were not recovered in any of the 21 contig bins compiled in this study, however, phylogenetic analyses using BLASTX to identify elongation factor G gene sequences with high % identities to the four contig bins and probable 16S rDNA OTUs revealed that the four bins selected for characterisation represented a Syntrophaceae species (bin 1.1), two *Syntrophorhabdus aromaticivorans* species (bins 2.2 and 2.6), and a novel member of the Fibrobacteres-Chlorobi-Bacteroidetes (FCB) superphylum (bin 3.1). Although the closest relative of bin 1.1 by its elongation factor G gene was a Syntrophaceae sp., possible Syntrophaceae matches for this bin in the 16S rDNA results from the metagenome were limited to two distinct taxa (OTU_64 and OTU_114; detected 12 and 10 times, respectively). Both of these have >90 % identities to type sequence *Smithella propionica* LYP (96.84 % and 94.86 %, respectively; GenBank accession NR_024989.1) as well as to each of the three formally described species within the *Syntrophus* genus (93.68–95.65 % and 92.89–94.09 %, respectively). Accordingly, bin 1.1 was referred to here as a Syntrophaceae sp., being closely related to the *Smithella* and *Syntrophus* genera. It is noteworthy that, based on phylogenomic analysis, there has been recent discussion around updating the phylogenetic ranks and assignments within the Deltaproteobacteria. Accordingly, *S. aromaticivorans* and Syntrophaceae sp. would be reclassified into a phylum named Thermodesulfobacteriota or Desulfobacterota, and, to incorporate high percent identities to both *S. propionica* and the *Syntrophus* spp., the nearest common rank would then be the Syntrophales order (42, 43) for bin 1.1. The nomenclature used in the present study does not reflect these proposed updates, and instead aligns with currently entries for Deltaproteobacteria and Syntrophaceae in the LPSN (https://lpsn.dsmz.de/).

Although 16S rDNA sequences were able to be matched by close relatives using only elongation factor G genes for the Deltaproteobacteria taxa (Syntrophaceae sp. bin 1.1, and *Syntrophorhabdus aromaticivorans* bins 2.2 and 2.6), the novelty of bin 3.1 impeded identification through close relatives.

Multilocus sequence analysis (MLSA) was performed to further characterise bin 3.1, after it was placed loosely within the FCB superphylum by low % identity NCBI BLAST results using its elongation factor G gene. MLSA was performed first on 37 representative genomes from the FCB superphylum (44), and then was also performed on 47 genomes from the Krumholzibacteriota, Delphibacteria, and Latescibacteria candidate phyla; the portion of the FCB superphylum that was most closely related to bin 3.1 (Supplementary Figure S2; Supplementary Table S4). These phyla are not yet well resolved, and the Genome Taxonomy Database classifies the Delphibacteria and some Latescibacteria within the Krumholzibacteriota phylum (45). From these MAGs, six housekeeping genes were selected for analysis: adenylosuccinate synthase, ATP synthase subunit beta, elongation factor G, elongation factor P, malate dehydrogenase, and protein RecA genes, which were identified in bin 3.1 in addition to a maximum number of the representative genomes. Importantly, each of these genes occurred within two or more genomes from Delphibacteria and Krumholzibacteriota; the two phyla with which bin 3.1 had the highest % identities in the BLASTX results for the elongation factor G gene. The results of the MLSA indicate that bin 3.1 is likely a novel member of the Krumholzibacteriota phylum (Figure 2), of which the closest known relative appears to be from a marine sediment enrichment culture from the Bothnian Sea (Supplementary Table S4) (46).

**Figure 2.**
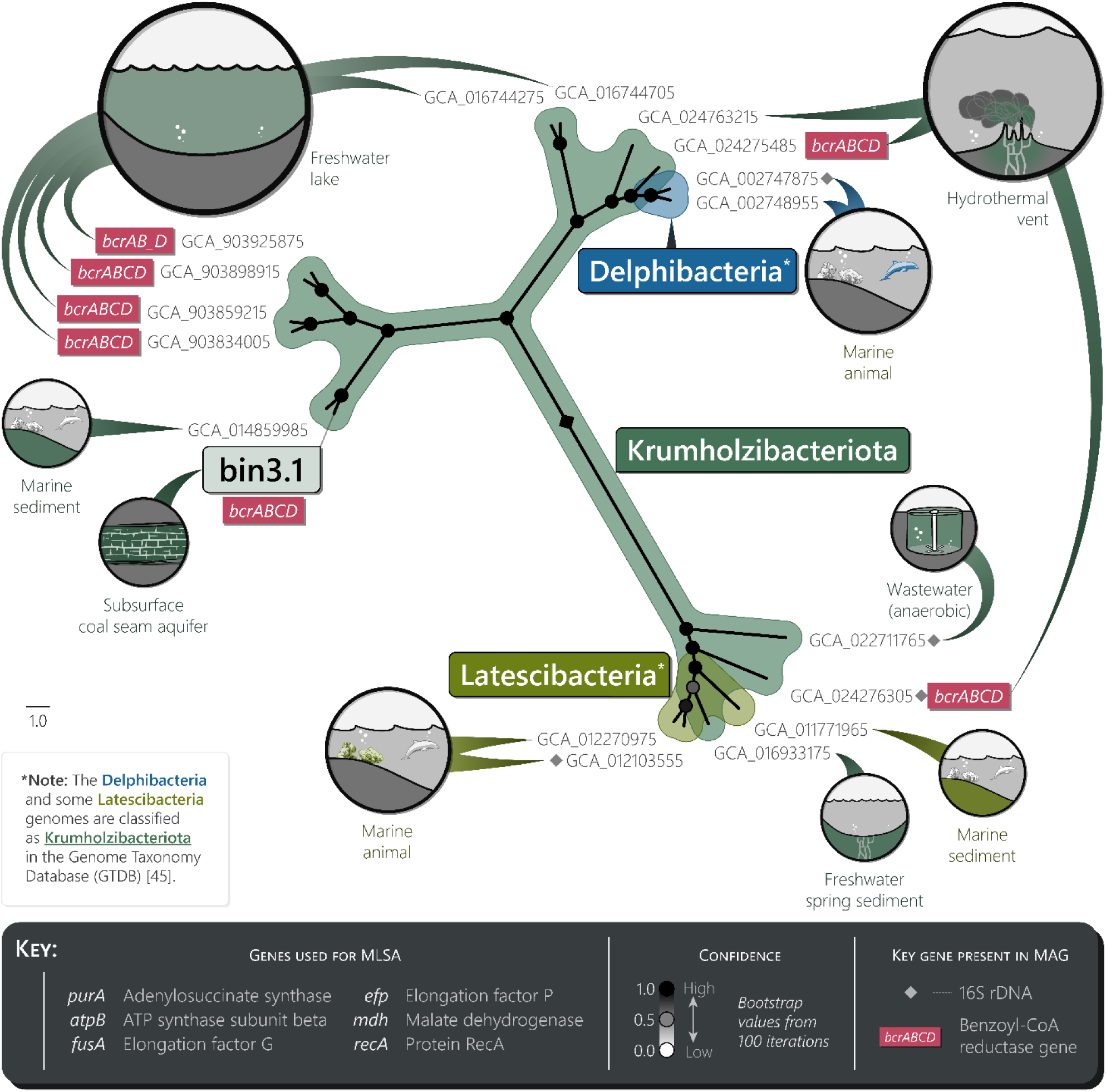
Phylogenetic relationships and environmental sources of selected Krumholzibacteriota, Delphibacteria*, and Latescibacteria* MAGs. Maximum likelihood phylogeny of bin 3.1 and 17 representative MAGs using multilocus sequence analysis (MLSA; see Supplementary Figure S2 for the equivalent FCB superphylum tree). Thirty other MAGs from the initial analysis were missing the selected housekeeping genes and thus are not included here (Supplementary Table S4) (46, 49, 50, 59–62). Environmental sources are indicated for each MAG; circle size is proportional to the number of times that environment applied within the same phylum.

For the purpose of aiding cross-study analysis of Krumholzibacteriota sp. bin 3.1, the four MAGs in the MLSA results that contained 16S rDNA sequences (Figure 2) were used to determine both the most probable 16S rDNA OTU in the present study, and the most probable CSMB reference set sequence (8). These four MAGs consisted of a Delphibacteria, two Krumholzibacteriota, and a Latescibacteria MAG (NCBI Assemblies GCA_002747875, GCA_022711765, GCA_024276305, and GCA_012103555, respectively). Each of these were compared against the metagenomic 16S rDNA OTUs and the CSMB reference set using USEARCH (90 % minimum threshold). Although no 16S rDNA OTU or CSMB identities were found for Krumholzibacteriota GCA_024276305 or Latescibacteria GCA_012103555, the USEARCH results indicated that Krumholzibacteriota sp. Bin 3.1 may correspond to OTU_57 (94.1 and 92.9 % identities; GCA_022711765 and GCA_002747875), which also is in agreement with the distribution of similar bin 3 group MAGs across the metagenomes in the present study (Supplementary Table S3; assuming loss of 16S rDNA sequences from rarer taxa in the more shallowly sequenced metagenomes). The CSMB identities indicated that Krumholzibacteriota sp. bin 3.1 may correspond to CSMB_1092 (95.5 and 93.9 % identities; GCA_022711765 and GCA_002747875), which has been previously reported from coal seams in the Bowen, Surat, and Sydney basins, Australia (Supplementary Table S5).

### General description of the Syntrophaceae sp., *S*. *aromaticivorans*, and Krumholzibacteriota sp. genome bins

The four bins chosen for further characterisation ranged in size from 1.96 to 4.18 Mbp, with *S. aromaticivorans* bin 2.6 having the smallest bin size, and the Krumholzibacteriota sp. having the largest bin size (Table 1). GC-content was broadly similar for the three Deltaproteobacteria MAGs (bins 1.1, 2.2 and 2.6), with an average of 46.7 %. Krumholzibacteriota sp. bin 3.1, however, had a higher GC-content of 71.1 % (Table 1) consistent with the other closely related Krumholzibacteriota MAGs from freshwater lakes and marine sediment (69.2 to 70.8 %; Figure 2; Supplementary Table S4) (46). Across the four bins, CheckM results indicated that contamination was very low (<2 %). Although only taxonomically resolved to the phylum level, genome completeness was greatest for Krumholzibacteriota sp. bin 3.1 (95 %), whilst genome completeness was lowest for the Powder River Basin-sourced *S. aromaticivorans* bin 2.6 (55 %; Table 1; Supplementary Table S3). Overall, these bins are medium- to high-quality draft MAGs, based on the combination of completeness and contamination scores with the varying presence of key marker genes within the assemblies (47). Although completeness and contamination scores for the Krumholzibacteriota sp. bin 3.1 were very good, it is ineligible for the classification of high-quality draft due to the lack of recovered rDNA sequences.

### Dearomatisation genes and other notable ecophysiological genomic characteristics

The most commonly detected benzoyl-CoA reductase gene subunits were those of *bcrABCD*, from the class-I enzyme system, associated with facultative anaerobes such as *Magnetospirillum* and *Thauera* spp. (15, 48). The Prokka annotations contained the highest number of these, however, the BlastKOALA annotations included an additional enzyme system (Table 2; Supplementary Table S2). This additional enzyme system, found in *S. aromaticivorans* bin 2.2, was the ATP-independent class-II benzoyl-CoA reductase gene subunit, *bamB*, associated with obligate anaerobes (Figure 1) (48).

**Table 2:**
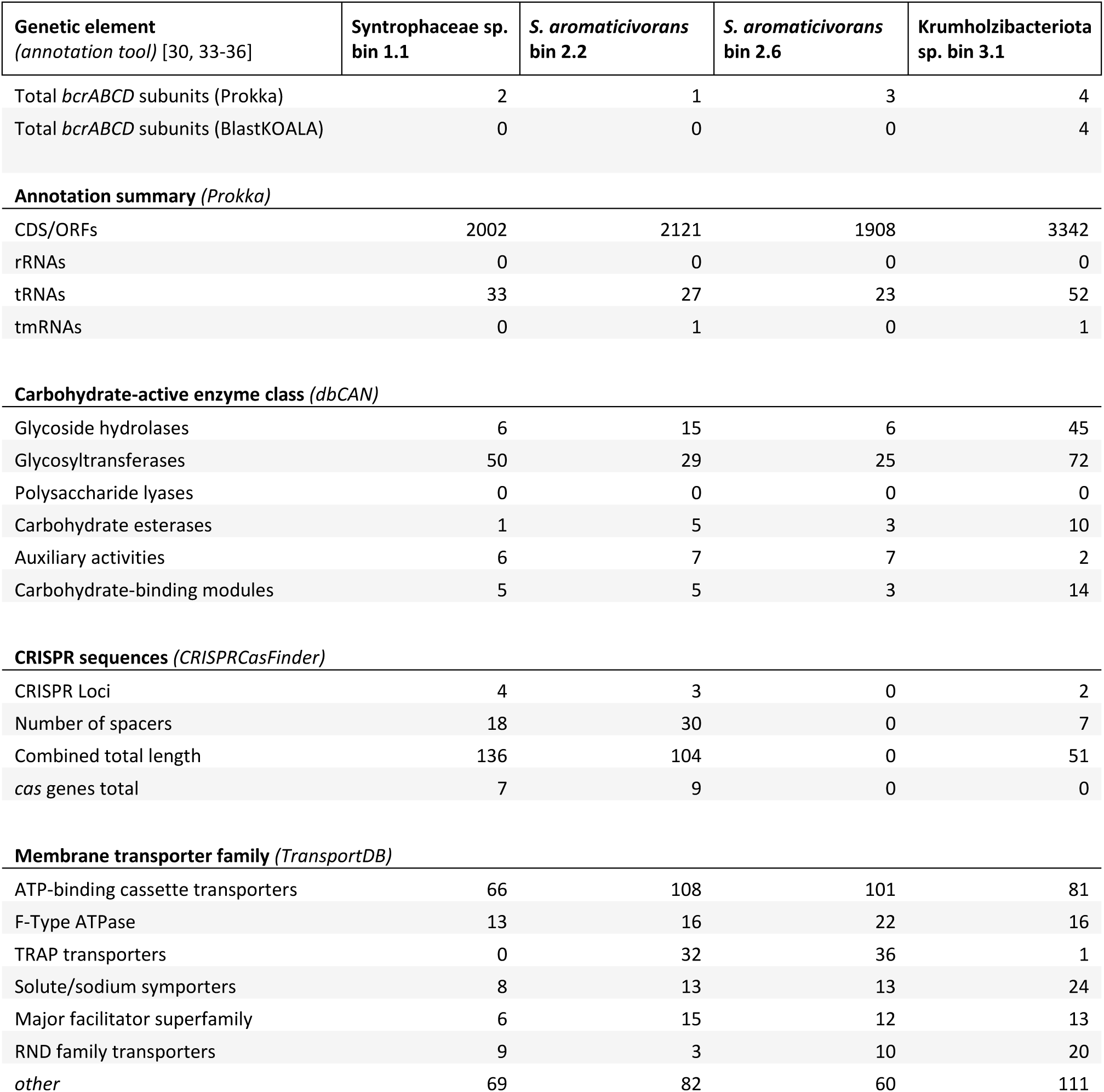
T**h**e **abundance of genes or genetic elements identified in the MAGs.**

In addition to their use for MLSA, the Prokka annotations of the representative MAGs from the FCB superphylum, including those from the Krumholzibacteriota, Latescibacteria, and Delphibacteria phyla, were searched for benzoyl-CoA pathway genes. From the 37 MAGs used for MLSA from the FCB superphylum, no genes of either the *bcr* or *bam* enzyme systems were detected (Supplementary Data Table S4a). Other genes from the benzoyl-CoA pathway were also rare, with the highest number (two) being found in the MAG of *Longimicrobium terrae* CECT 8660, from the Gemmatimonadetes phylum, for the production of benzoyl-CoA from 4-hydroxybenzoyl-CoA and benzoate (*hcrABC* and *badA*; respectively). In contrast, the *bcr* gene and other benzoyl-CoA reductase genes were far more common among the Krumholzibacteriota, Latescibacteria, and Delphibacteria MAGs used for MLSA: 17 out of 40 contained one or more subunits from *bcrABCD*, with 12 of these containing all four subunits (Supplementary Data Table S4b) (46, 49, 50). Further, 25 of the 40 MAGs contained one or more genes (or gene subunits) from the benzoyl-CoA pathway. Of these, four taxa contained an identical distribution of pathway genes to Krumholzibacteriota sp. bin 3.1, and were sourced from a hydrothermal vent chimney, and freshwater lakes in Puerto Rico and Finland (GCA_024275485, GCA_903859215, GCA_903912525, and GCA_903834005).

Aside from the benzoyl-CoA pathway genes, the most notable BlastKOALA results regarded flagellar assembly and chemotaxis, anabolic antimicrobial genes, and nutrient and electron acceptor scavenging. Substantial numbers of genes for flagellar assembly and chemotaxis were detected within Krumholzibacteriota sp. bin 3.1, as well as genes for biosynthesis of antimicrobial substances. Genes for nutrient and electron acceptor scavenging of nitrogen and sulfur compounds were most commonly detected within the *S. aromaticivorans* bins 2.2 and 2.6.

### Carbohydrate-active enzymes

Each of the four bins contained a small number of carbohydrate-active enzymes (Table 2; Supplementary Table S2). The highest number of carbohydrate-active enzymes was detected in Krumholzibacteriota sp. bin 3.1 (143 total), of which the majority were glycosyltransferases and glycoside hydrolases. The lowest number of carbohydrate-active enzymes was found in *S. aromaticivorans* bin 2.6 (44 total), more than half of which were glycosyltransferases. Overall, glycosyltransferases were the most commonly detected carbohydrate-active enzymes in each bin. No polysaccharide lyases were detected in any of the bins.

### Membrane transporter proteins

Numerous transporter protein genes were detected in each bin (Table 2; Supplementary Table S2). *Syntrophorhabdus aromaticivorans* bins 2.2 and 2.6, along with Krumholzibacteriota sp. bin 3.1, contained higher numbers of transporter proteins (totalling 269, 254 and 266 respectively). In contrast, Syntrophaceae sp. bin 1.1 contained a lower number of transporter proteins (totalling 171). Across all four bins, the ATP-binding cassette transporters were the most commonly detected, however, it is also notable that the tripartite ATP-independent periplasmic transporters were unusually abundant within the *S. aromaticivorans* bins 2.2 and 2.6 (32 and 36, respectively).

### CRISPRs and *cas* genes

Three of the four bins included multiple CRISPR loci, and two of these (bins 1.1 and 2.2) contained several *cas* genes (Table 2; Supplementary Table S2). Syntrophaceae sp. bin 1.1 had four CRISPR loci containing 18 spacers, *S. aromaticivorans* bin 2.2 had three CRISPR loci and 30 spacers, and Krumholzibacteriota sp. bin 3.1 had two CRISPR loci and seven spacers. Notably, while Syntrophaceae sp. bin 1.1 and *S. aromaticivorans* bin 2.2 contained the aforementioned *cas* genes (7 and 9, respectively), no *cas* genes were detected within Krumholzibacteriota sp. bin 3.1 and neither CRISPR loci nor *cas* genes were detected in *S. aromaticivorans* bin 2.6.

## DISCUSSION

For microbiologists working to understand the catabolism of coal in the subsurface, the identification of taxa engaged in degrading aromatic compounds has proved elusive. Indeed, microbial communities from the coal seam environment tend to be numerically dominated by one or two methanogens (12, 51), a handful of bacterial taxa presumably engaging in syntrophic partnerships with these methanogens, and a long tail of rarer taxa (52). It has generally been the view that bacterial syntrophs could not degrade complex aromatic or aliphatic compounds (4, 53, 54), which leaves the largely unexplored tail of the microbial community in coal as a likely place to find those microbes capable of coal degradation. Further, it is ecologically reasonable to hypothesise that this unexplored tail contains microbes with a more diverse array of nutritional strategies, since presumably there are abundant and less-highly competed nutritional niches within the diverse heterogeneity of the organic matter in coal. Data presented here demonstrate for the first time the identity of multiple lineages with crucial monoaromatic-degrading genes from subsurface coal seams on two continents. Four taxa were observed in this study: one Syntrophaceae sp. (bin 1.1), two *Syntrophorhabdus aromaticivorans* (bins 2.2 and 2.6) and a novel taxon from the Krumholzibacteriota phylum in the FCB superphylum (bin 3.1). This Krumholzibacteriota sp. is likely engaged in monoaromatic degradation in the Powder River Basin, and had not previously been implicated as having a role in these processes in the coal seam environment. While it has been previously stated that *S. aromaticivorans* may be important for coal degradation (22), this is the first demonstration that coal seam strains of this taxon carry genes for monoaromatic degradation. For the Syntrophaceae sp., taxa within this family are well-known hydrocarbon-degraders in subsurface environments, such as aliphatic molecule degradation in oil reservoirs (26), however, they had not previously been implicated in aromatic degradation in the coal seam environment.

### Genomic characterisation of the Krumholzibacteriota sp. bin3.1 MAG

Krumholzibacteriota taxa are identified here for the first time as aromatic-degraders in the coal seam environment, as well as more broadly. From the annotations of its genome, the Krumholzibacteriota sp. is a primary coal degrader with the ability to anaerobically catabolise a wide range of monoaromatic compounds (Figure 3; Supplementary Table S2). Further, the Krumholzibacteriota sp. may also compete in other ecological niches such as biomass recycling, since in addition to genes for monoaromatic degradation, it also has a high number of glycoside hydrolase genes (relative to the three Deltaproteobacteria MAGs; Table 2). It is probable that the organism is performing biomass recycling to supplement its catabolism of carbon from aromatic substrates, rather than specialising in biomass recycling, since the abundance of glycoside hydrolase genes is lower in the Krumholzibacteriota sp. relative to other biomass degraders described from the coal seam environment (52). In addition to being able to obtain nitrogen and phosphorus through biomass recycling, the Krumholzibacteriota sp. may be capable of extracting nutrients from coal via the degradation of nitrogen and phosphate-containing aromatic compounds, since it has a near-complete set of genes for the catabolism of benzamide and benzoyl phosphate to acetate (Figure 3). Moreover, it is also the only MAG described here to contain most of the genes necessary for flagellar assembly and chemotaxis, which it may use to seek out desirable regions of organic matter within the coal. Finally, the Krumholzibacteriota sp. contained a high number of multidrug efflux pumps relative to the other organisms examined here, which may be relevant for removal of toxic metabolites produced during catabolism of aromatic substrates. Overall, these findings implicate this Krumholzibacteriota sp. as potentially important for both the dearomatisation of monoaromatic compounds, and the production of simpler methanogenic substrates in the coal seam environment. Further studies verifying the function of this microbe in the coal seam environment are needed, ideally in the form of axenic culturing to test its response to different variables such as complex carbon substrates. Axenic culturing would also provide details on the cell morphology of this Krumholzibacteriota sp., and may aid in understanding its ability to access different regions within the coal seam environment. Such efforts to obtain this taxon in pure culture may be tailored using its described characteristics from the genome provided here.

**Figure 3.**
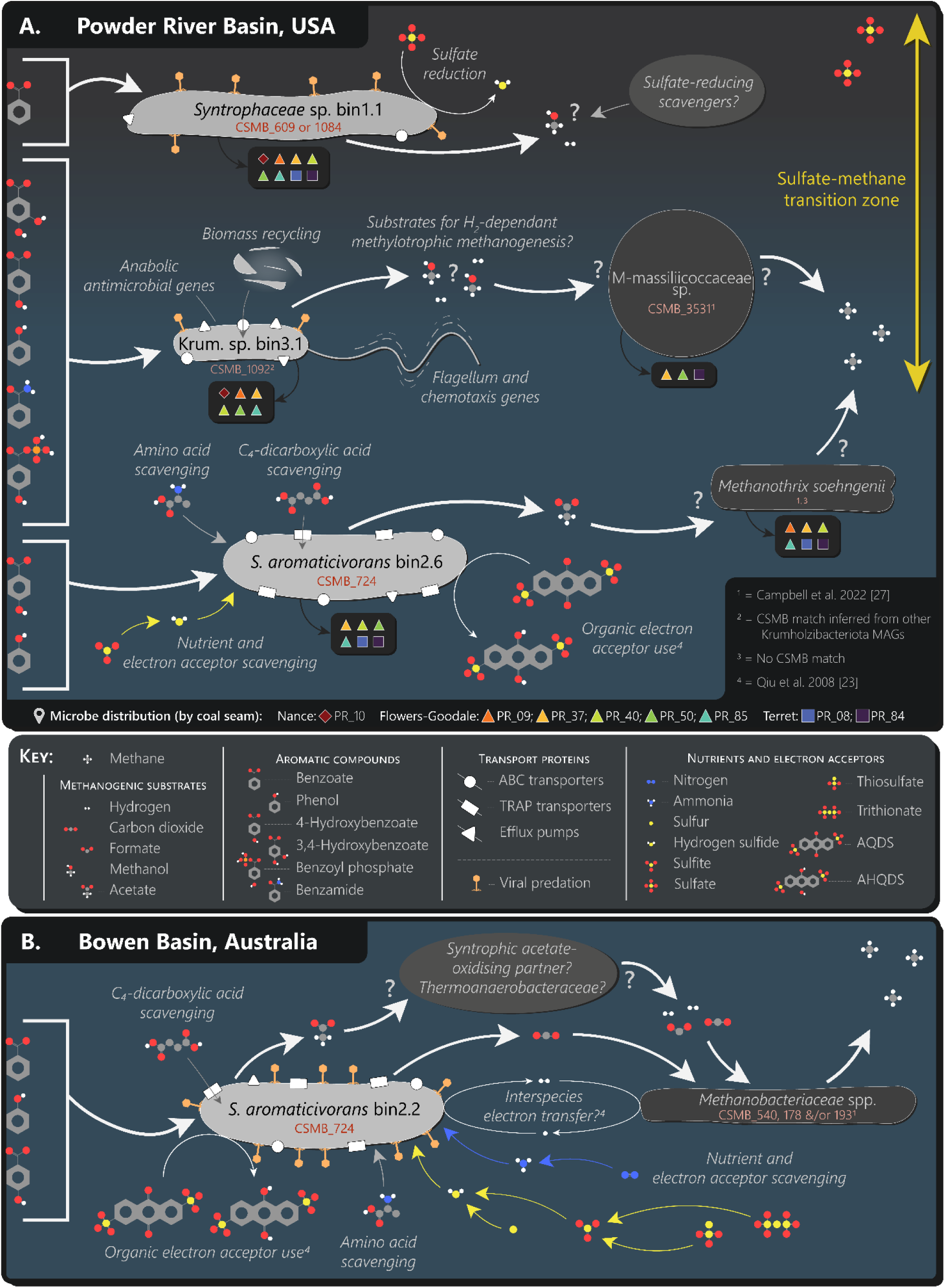
Putative ecological functions within the coal seam microbiome. Conceptual model displaying (A) the Syntrophaceae sp., Krumholzibacteriota sp. and *Syntrophorhabdus aromaticivorans* within the Nance, Flowers-Goodale and Terret coal seams of the Powder River Basin, USA, and (B) the *Syntrophorhabdus aromaticivorans* within the Bandanna Formation of the Bowen Basin, Australia. The most abundant methanogen/s (according to *mcrA* and 16S rDNA data; (27)) in the metagenome of each bin has been included in the model, however, these are not necessarily the key partner methanogens for the aromatic-degrading taxa.

Taxonomically, the classification of this Krumholzibacteriota sp. was more challenging than for the Deltaproteobacteria MAGs, both because the assembly contained no annotated sequences of 16S rDNA, and because the elongation factor G gene did not return any high % identities to other well characterised genomes in the NCBI collection. The results of the MLSA indicate that the closest relatives of this Krumholzibacteriota sp. were other taxa from within the same phylum, found in marine sediments from the Bothnian Sea in Scandinavia (46), a stratified freshwater reservoir of the Cañas River in Puerto Rico, and Lake Lovojärvi in Finland (Figure 2; Supplementary Table S4). Although 16S rDNA sequences for the Krumholzibacteriota sp. (OTU_57; CSMB_1092) were able to be inferred using four representative MAGs of the same phylum, confirmation of the 16S rDNA sequences specific to this Krumholzibacteriota sp. is needed for global context in the subsurface coal seam environment, as well as other environments where the Krumholzibacteriota spp. may play a key ecological role.

### Krumholzibacteriota is a globally-relevant phylum for liberation of recalcitrant carbon

Indeed, dearomatisation and downstream catabolism of monoaromatic compounds in anoxic environments may be relatively common within the Krumholzibacteriota phylum (Figure 2). Of the 40 other Krumholzibacteriota MAGs used for MLSA, 13 contained a complete or near-complete set of genes for catabolism of benzoyl-CoA to 3-hydroxypimeloyl-CoA (Supplementary Table S4). In addition to this genetic potential for anaerobic dearomatisation, aerobic dearomatisation has recently been suggested regarding a Krumholzibacteriota MAG that contained one subunit of the gene encoding for benzoate/toluate 1,2-dioxygenase reductase (*benC*-*xylZ*), and was sourced from an oil-contaminated environment in the Persian Gulf (55). Again, axenic studies of these carbon-liberating Krumholzibacteriota would be beneficial in clarifying the phylogenetic and metabolic capabilities of taxa within this phylum. Specifically, this clarification is essential for a clearer understanding of the potential rate-limiting ecological roles of Krumholzibacteriota spp. in the subsurface coal seam environment. Axenic studies can also aid in understanding the roles of these organisms in other carbon-cycling environments where they have been detected, such as marine sediments, deep marine cold seep fluids, hydrothermal vents, and surface freshwaters.

### Syntrophorhabdus aromaticivorans in the coal seam environment

Unlike the novel Krumholzibacteriota sp., *S. aromaticivorans* has been identified within the coal seam environment on numerous previous occasions. Indeed, the CSMB reference set presently lists six distinct taxa from the *Syntrophorhabdus* genus in coal-bearing basins of Australia (Supplementary Table S5), and they have also been detected in the Powder River Basin in the USA (22). A recent study of microbial communities from the Powder River Basin implicated this taxon as an important potential primary degrader of organic matter in coal, based on both its increases in abundance after algal-amendment, and the known degradative abilities of the single strain of this taxon in culture (23). It should be noted, however, that only a single isolate of this species has been described, and this isolate was obtained from wastewater, not from the subsurface. It was thus unclear whether other strains of *S. aromaticivorans* were capable of aromatic degradation.

Data from the current study demonstrate, for the first time, that two other genomes of *S. aromaticivorans* have at least partial pathways for the degradation of monoaromatic compounds (Figure 1). This is likely to be a common feature in the genome of this species, given that these two new strains come from different coal seams on different continents. Taken together, these data indicate that *S. aromaticivorans* can likely directly access carbon contained within coal via aromatic molecules, such as 4-hydroxybenzoate and benzoate. In addition to its ability to catabolise monoaromatic compounds, both the North American and Australian *S. aromaticivorans* strains described here appear to have an unusually large array of tripartite ATP-independent periplasmic (TRAP) transporters, which may indicate that dicarboxylates (such as fumarate) are an important source of carbon for *S. aromaticivorans* (Figure 3). Biomass recycling, on the other hand, is unlikely to be a source of carbon for this taxon, as neither strain contained substantial numbers of genes for glycoside hydrolase and other carbohydrate enzymes. Interestingly, the Australian *S. aromaticivorans* genome contains abundant CRISPR spacers, suggesting substantial pressure on this taxon from viral predation in the Bowen Basin. In contrast, the lack of CRISPR sequences detected in the North American genome indicates little to no viral predation stress on this taxon in the Powder River Basin. Viral predation has been previously implicated as an important process in the Surat Basin (14), and numerous other taxa from this environment have been previously demonstrated to harbour significantly CRISPR spacers arrayed against a host of viruses (10, 56). The absence of these sequences in the Powder River Basin may indicate this species, at least, is not under the same level of viral predation as in other subsurface systems.

Alternatively, it may be that these sequences were simply not recovered due to the insufficient completeness of this bin (Table 1). Lastly, the ability of *S. aromaticivorans* to use organic electron acceptors (9,10-anthraquinone-2,6-disulfonate) when isolated from other environments is intriguing (23), and it may be that aromatic compounds from the coal seam environment could also act as alternative electron acceptors, though experimental work with the organism in culture would be important to confirm this speculation. Regardless, certainly for Australian strains, there is clear evidence that this species has the genetic potential to catabolise aromatic compounds from the coal seam, and, if these taxa are also subject to viral predation, it may be a mechanism by which this carbon is made available to a wider range of taxa post-lysis of the *S. aromaticivorans* cells.

### Syntrophaceae taxa may utilise aromatic carbon in coal seams

The other Deltaproteobacteria taxon identified was a bin associated with the family Syntrophaceae, having genes for monoaromatic degradation in the Powder River Basin. Data from the CSMB reference set reveal that the Syntrophaceae family is commonly present within coal seams in Australia, in all of the North American coal seams in the present study, and also in the Ishikari Basin, Japan (Supplementary Table S5; (8)). The Syntrophaceae family, and *Smithella propionica* specifically, is well known for its use of aliphatic hydrocarbons (26, 57). Indeed, *Smithella* spp. are well-known alkane-degraders from anoxic environments such as oil reservoirs, where they can grow in syntrophy with hydrogenotrophic methanogens (26). Despite this prevalence in hydrocarbon-rich environments, Syntrophaceae sp. bin 1.1 represents the first time that this family has been implicated in primary degradation of aromatic substrates in the coal seam environment. Of additional note regarding metabolism and ecological niches of this taxon, is the absence of substantial numbers of genes for carbohydrate degradation. This absence, in addition to its low number of ABC transporters, suggests that it does not participate in biomass recycling, nor does it experience substantial competitive stress. This genome does, however, appear to experience considerable pressure from viral predation, as it contains a high number of CRISPR spacers relative to the other Powder River Basin taxa examined here (Figure 3; Table 2). The Syntrophaceae sp. co-occurs with the relatively viral predation-free *S. aromaticivorans* in the Powder River Basin, and it thus seems likely that viral predation may be an important process in the Powder River Basin, as it is in other subsurface habitats.

### Ecological functions and implications for understanding the coal seam microbiome

In terms of their life strategies in the coal seam, all three taxa likely have stress-tolerant ecological profiles (in the sense of Grime (58)). All three appear to possess specialised genetic tools for the degradation of a plentiful, but difficult to access, carbon source within subsurface coal seams. Interestingly, the Krumholzibacteriota taxon, as well as hosting these genes for monoaromatic degradation, also hosts an array of other genes for accessing nutrients in moribund cells or plant material (Figure 3; Table 2). As described earlier, coal seams are oligotrophic environments, and access to other macronutrients, such as nitrogen, in coal organic matter may be important for competition in this environment. Further, while it appears that the Krumholzibacteriota organism described here may have a relatively high number of genes involved in carbohydrate metabolism, it has relatively few compared to truly ruderal taxa that occur in coal seams (14).

Importantly for scientific efforts to enhance or control methane yields from coal, the four putative aromatic-degrading taxa described here may hold key rate-limiting roles in the biodegradation of coal to methane in the subsurface. If the Krumholzibacteriota sp., the two *S. aromaticivorans*, and the Syntrophaceae sp. are all indeed capable of direct access to the carbon within coal, further study of their metabolic strategies may provide important tools for altering biodegradation of coal in the subsurface. One key goal of future research should be to obtain isolates of each of these taxa, which would not only be beneficial in confirming their function, but also provide a suite of tools for further manipulative experiments.

Much of the previous research into the coal seam microbiome has centred around either descriptive studies of species distributions, or the effect of nutrient mixtures in an effort to enhance gas yields. While both of these approaches are valuable, they provide comparatively little information on the function of individual microbes in these communities. In contrast, this study describes the genomes of four new taxa from the coal seam environment with likely roles in monoaromatic compound degradation. Studies that seek to elucidate processes upstream of monoaromatic degradation, involving the liberation of soluble organic matter from the insoluble coal macromolecule, would further our understanding of this unusual environment.

## Acknowledgements

The authors thank Dr. Andrew G. McLeish for running the contig bins with CheckM. Dr. Bronwyn Campbell was supported by a Macquarie University Research Training Program scholarship, a CSIRO Energy strategic research initiative, and a PESA Horstmann Federal Postgraduate Scholarship.

## Competing interests statement

The authors declare no competing interests.

## Data availability

The datasets sourced for analysis in the current study were downloaded from the NCBI Sequence Read Archive (from BioProject accessions PRJNA330673, PRJNA291107, and PRJNA678021; sequence run accessions can be found in Supplementary Table S1), the JGI Data Portal (Gold project IDs Gp0406113 to Gp0406117, and the CSIRO Data Access Portal (https://doi.org/10.4225/08/5b31ca6373d48).

The MAGs (as contig bins and Prokka annotations), 16S rDNA OTU sequences, and CSMB-OTU 16S rDNA data table generated during the current study are each available in the CSIRO Data Access Portal (https://doi.org/10.25919/d4nf-8f32).

## Notes

### Competing Interest Statement

The authors have declared no competing interest.

### Summary of Updates

Removal of an author from the author list until full review process is completed by USGS.

## Reference List

1. Hardisty PE, Clark TS, Hynes RG. 2012. Life Cycle Greenhouse Gas Emissions from Electricity Generation: A Comparative Analysis of Australian Energy Sources. 4. Energies 5:872–897.

2. Markandya A, Wilkinson P. 2007. Electricity generation and health. The Lancet 370:979–990.

3. Schandl H, Baynes T, Haque N, Barrett D, Geschke A. 2019. Final Report for GISERA Project G2 - Whole of Life Greenhouse Gas Emissions Assessment of a Coal Seam Gas to Liquefied Natural Gas Project in the Surat Basin, Queensland, Australia. CSIRO Aust.

4. Strąpoć D, Mastalerz M, Dawson K, Macalady J, Callaghan AV, Wawrik B, Turich C, Ashby M. 2011. Biogeochemistry of Microbial Coal-Bed Methane. Annu Rev Earth Planet Sci 39:617–656.

5. Ritter D, Vinson D, Barnhart E, Akob DM, Fields MW, Cunningham AB, Orem W, McIntosh JC. 2015. Enhanced microbial coalbed methane generation: A review of research, commercial activity, and remaining challenges. Int J Coal Geol 146:28–41.

6. Wang H, Lin H, Rosewarne CP, Li D, Gong S, Hendry P, Greenfield P, Sherwood N, Midgley DJ. 2016. Enhancing biogenic methane generation from a brown coal by combining different microbial communities. Int J Coal Geol 154–155:107–110.

7. He X, Liu X, Nie B, Song D. 2017. FTIR and Raman spectroscopy characterization of functional groups in various rank coals. Fuel 206:555–563.

8. Vick SHW, Greenfield P, Tran-Dinh N, Tetu SG, Midgley DJ, Paulsen IT. 2018. The Coal Seam Microbiome (CSMB) reference set, a *lingua franca* for the microbial coal-to-methane community. Int J Coal Geol 186:41–50.

9. Wang B, Yu Z, Zhang Y, Zhang H. 2019. Microbial communities from the Huaibei Coalfield alter the physicochemical properties of coal in methanogenic bioconversion. Int J Coal Geol 202:85–94.

10. McLeish AG, Greenfield P, Midgley DJ, Paulsen IT. 2021. Draft Genome Sequence of *Desulfovibrio* sp. Strain CSMB_222, Isolated from Coal Seam Formation Water. Microbiol Resour Announc https://doi.org/10.1128/MRA.00564-21.

11. Johnson ER, Klasson KT, Basu R, Volkwein JC, Clausen EC, Gaddy JL. 1994. Microbial conversion of high-rank coals to methane. Appl Biochem Biotechnol 45:329.

12. McKay LJ, Smith HJ, Barnhart EP, Schweitzer HD, Malmstrom RR, Goudeau D, Fields MW. 2021. Activity-based, genome-resolved metagenomics uncovers key populations and pathways involved in subsurface conversions of coal to methane. ISME J 1–12.

13. Schweitzer HD, Smith HJ, Barnhart EP, McKay LJ, Gerlach R, Cunningham AB, Malmstrom RR, Goudeau D, Fields MW. 2022. Subsurface hydrocarbon degradation strategies in low- and high-sulfate coal seam communities identified with activity-based metagenomics. 1. Npj Biofilms Microbiomes 8:1–10.

14. Vick SHW, Greenfield P, Tetu SG, Midgley DJ, Paulsen IT. 2019. Genomic and phenotypic insights point to diverse ecological strategies by facultative anaerobes obtained from subsurface coal seams. 1. Sci Rep 9:1–13.

15. Carmona M, Zamarro MT, Blázquez B, Durante-Rodríguez G, Juárez JF, Valderrama JA, Barragán MJL, García JL, Díaz E. 2009. Anaerobic catabolism of aromatic compounds: a genetic and genomic view. Microbiol Mol Biol Rev MMBR 73:71–133.

16. Foght J. 2008. Anaerobic Biodegradation of Aromatic Hydrocarbons: Pathways and Prospects. Microb Physiol 15:93–120.

17. Boll M, Fuchs G. 1995. Benzoyl-Coenzyme A Reductase (Dearomatizing), a Key Enzyme of Anaerobic Aromatic Metabolism. Eur J Biochem 234:921–933.

18. McLeish AG, Gong S, Greenfield P, Midgley DJ, Paulsen IT. 2021. Microbial Community Shifts on Organic Rocks of Different Maturities Reveal potential Catabolisers of Organic Matter in Coal. Microb Ecol https://doi.org/10.1007/s00248-021-01857-x.

19. Campbell BC, Gong S, Greenfield P, Midgley DJ, Paulsen IT, George SC. 2021. Aromatic compound-degrading taxa in an anoxic coal seam microbiome from the Surat Basin, Australia. FEMS Microbiol Ecol 97:fiab053.

20. Kühner S, Wöhlbrand L, Fritz I, Wruck W, Hultschig C, Hufnagel P, Kube M, Reinhardt R, Rabus R. 2005. Substrate-Dependent Regulation of Anaerobic Degradation Pathways for Toluene and Ethylbenzene in a Denitrifying Bacterium, Strain EbN1. J Bacteriol 187:1493–1503.

21. Schink B, Philipp B, Müller J. 2000. Anaerobic degradation of phenolic compounds. Naturwissenschaften 87:12–23.

22. Smith HJ, Schweitzer HD, Barnhart EP, Orem W, Gerlach R, Fields MW. 2021. Effect of an algal amendment on the microbial conversion of coal to methane at different sulfate concentrations from the Powder River Basin, USA. Int J Coal Geol 248:103860.

23. Qiu Y-L, Hanada S, Ohashi A, Harada H, Kamagata Y, Sekiguchi Y. 2008. Syntrophorhabdus aromaticivorans gen. nov., sp. nov., the First Cultured Anaerobe Capable of Degrading Phenol to Acetate in Obligate Syntrophic Associations with a Hydrogenotrophic Methanogen. Appl Environ Microbiol 74:2051–2058.

24. Vick SHW, Gong S, Sestak S, Vergara TJ, Pinetown KL, Li Z, Greenfield P, Tetu SG, Midgley DJ, Paulsen IT. 2019. Who eats what? Unravelling microbial conversion of coal to methane. FEMS Microbiol Ecol 95:fiz093.

25. Kuever J. 2014. The Family Syntrophaceae, p. 281–288. In Rosenberg, E, DeLong, EF, Lory, S, Stackebrandt, E, Thompson, F (eds.), The Prokaryotes: Deltaproteobacteria and Epsilonproteobacteria. Springer, Berlin, Heidelberg.

26. Gray ND, Sherry A, Grant RJ, Rowan AK, Hubert CRJ, Callbeck CM, Aitken CM, Jones DM, Adams JJ, Larter SR, Head IM. 2011. The quantitative significance of Syntrophaceae and syntrophic partnerships in methanogenic degradation of crude oil alkanes. Environ Microbiol 13:2957–2975.

27. Campbell BC, Greenfield P, Gong S, Barnhart EP, Midgley DJ, Paulsen IT, George SC. 2022. Methanogenic archaea in subsurface coal seams are biogeographically distinct: an analysis of metagenomically-derived mcrA sequences. Environ Microbiol 24:4065–4078.

28. Greenfield P, Duesing K, Papanicolaou A, Bauer DC. 2014. Blue: correcting sequencing errors using consensus and context. Bioinformatics 30:2723–2732.

29. Prjibelski A, Antipov D, Meleshko D, Lapidus A, Korobeynikov A. 2020. Using SPAdes De Novo Assembler. Curr Protoc Bioinforma 70:e102.

30. Seemann T. 2014. Prokka: rapid prokaryotic genome annotation. Bioinformatics 30:2068– 2069.

31. Dick GJ, Andersson AF, Baker BJ, Simmons SL, Thomas BC, Yelton AP, Banfield JF. 2009. Community-wide analysis of microbial genome sequence signatures. Genome Biol 10:R85.

32. Parks DH, Imelfort M, Skennerton CT, Hugenholtz P, Tyson GW. 2015. CheckM: assessing the quality of microbial genomes recovered from isolates, single cells, and metagenomes. Genome Res 25:1043–1055.

33. Kanehisa M, Sato Y, Morishima K. 2016. BlastKOALA and GhostKOALA: KEGG Tools for Functional Characterization of Genome and Metagenome Sequences. J Mol Biol 428:726– 731.

34. Elbourne LDH, Tetu SG, Hassan KA, Paulsen IT. 2017. TransportDB 2.0: a database for exploring membrane transporters in sequenced genomes from all domains of life. Nucleic Acids Res 45:D320–D324.

35. Zhang H, Yohe T, Huang L, Entwistle S, Wu P, Yang Z, Busk PK, Xu Y, Yin Y. 2018. dbCAN2: a meta server for automated carbohydrate-active enzyme annotation. Nucleic Acids Res 46:W95–W101.

36. Couvin D, Bernheim A, Toffano-Nioche C, Touchon M, Michalik J, Néron B, Rocha EPC, Vergnaud G, Gautheret D, Pourcel C. 2018. CRISPRCasFinder, an update of CRISRFinder, includes a portable version, enhanced performance and integrates search for Cas proteins. Nucleic Acids Res 46:W246–W251.

37. Greenfield P, Tran-Dinh N, Midgley D. 2019. Kelpie: generating full-length ‘amplicons’ from whole-metagenome datasets. PeerJ 6:e6174.

38. Apprill A, McNally S, Parsons R, Weber L. 2015. Minor revision to V4 region SSU rRNA 806R gene primer greatly increases detection of SAR11 bacterioplankton. Aquat Microb Ecol 75:129–137.

39. Parada AE, Needham DM, Fuhrman JA. 2016. Every base matters: assessing small subunit rRNA primers for marine microbiomes with mock communities, time series and global field samples. Environ Microbiol 18:1403–1414.

40. Edgar RC. 2010. Search and clustering orders of magnitude faster than BLAST. Bioinformatics 26:2460–2461.

41. Tamura K, Stecher G, Kumar S. 2021. MEGA11: Molecular Evolutionary Genetics Analysis Version 11. Mol Biol Evol 38:3022–3027.

42. Waite DW, Chuvochina M, Pelikan C, Parks DH, Yilmaz P, Wagner M, Loy A, Naganuma T, Nakai R, Whitman WB, Hahn MW, Kuever J, Hugenholtz P. 2020. Proposal to reclassify the proteobacterial classes Deltaproteobacteria and Oligoflexia, and the phylum Thermodesulfobacteria into four phyla reflecting major functional capabilities. Int J Syst Evol Microbiol 70:5972–6016.

43. Oren A, Garrity GM. 2021. Valid publication of the names of forty-two phyla of prokaryotes. Int J Syst Evol Microbiol 71:005056.

44. Hug LA, Baker BJ, Anantharaman K, Brown CT, Probst AJ, Castelle CJ, Butterfield CN, Hernsdorf AW, Amano Y, Ise K, Suzuki Y, Dudek N, Relman DA, Finstad KM, Amundson R, Thomas BC, Banfield JF. 2016. A new view of the tree of life. 5. Nat Microbiol 1:1–6.

45. Parks DH, Chuvochina M, Rinke C, Mussig AJ, Chaumeil P-A, Hugenholtz P. 2022. GTDB: an ongoing census of bacterial and archaeal diversity through a phylogenetically consistent, rank normalized and complete genome-based taxonomy. Nucleic Acids Res 50:D785–D794.

46. Dalcin Martins P, de Jong A, Lenstra WK, van Helmond NAGM, Slomp CP, Jetten MSM, Welte CU, Rasigraf O. 2021. Enrichment of novel Verrucomicrobia, Bacteroidetes, and Krumholzibacteria in an oxygen-limited methane- and iron-fed bioreactor inoculated with Bothnian Sea sediments. MicrobiologyOpen 10:e1175.

47. Bowers RM, Kyrpides NC, Stepanauskas R, Harmon-Smith M, Doud D, Reddy TBK, Schulz F, Jarett J, Rivers AR, Eloe-Fadrosh EA, Tringe SG, Ivanova NN, Copeland A, Clum A, Becraft ED, Malmstrom RR, Birren B, Podar M, Bork P, Weinstock GM, Garrity GM, Dodsworth JA, Yooseph S, Sutton G, Glöckner FO, Gilbert JA, Nelson WC, Hallam SJ, Jungbluth SP, Ettema TJG, Tighe S, Konstantinidis KT, Liu W-T, Baker BJ, Rattei T, Eisen JA, Hedlund B, McMahon KD, Fierer N, Knight R, Finn R, Cochrane G, Karsch-Mizrachi I, Tyson GW, Rinke C, Lapidus A, Meyer F, Yilmaz P, Parks DH, Murat Eren A, Schriml L, Banfield JF, Hugenholtz P, Woyke T. 2017. Minimum information about a single amplified genome (MISAG) and a metagenome-assembled genome (MIMAG) of bacteria and archaea. 8. Nat Biotechnol 35:725–731.

48. von Netzer F, Granitsiotis MS, Szalay AR, Lueders T. 2020. Next-Generation Sequencing of Functional Marker Genes for Anaerobic Degraders of Petroleum Hydrocarbons in Contaminated Environments, p. 257–276. In Boll, M (ed.), Anaerobic Utilization of Hydrocarbons, Oils, and Lipids. Springer International Publishing, Cham.

49. Murphy CL, Biggerstaff J, Eichhorn A, Ewing E, Shahan R, Soriano D, Stewart S, VanMol K, Walker R, Walters P, Elshahed MS, Youssef NH. 2021. Genomic characterization of three novel Desulfobacterota classes expand the metabolic and phylogenetic diversity of the phylum. Environ Microbiol 23:4326–4343.

50. Vega MAP, Scholes RC, Brady AR, Daly RA, Narrowe AB, Bosworth LB, Wrighton KC, Sedlak DL, Sharp JO. 2022. Pharmaceutical Biotransformation is Influenced by Photosynthesis and Microbial Nitrogen Cycling in a Benthic Wetland Biomat. Environ Sci Technol 56:14462–14477.

51. Rosewarne CP, Greenfield P, Li D, Tran-Dinh N, Midgley DJ, Hendry P. 2013. Draft Genome Sequence of *Methanobacterium* sp. Maddingley, Reconstructed from Metagenomic Sequencing of a Methanogenic Microbial Consortium Enriched from Coal-Seam Gas Formation Water. Genome Announc 1:e00082–12.

52. Vick SHW, Greenfield P, Pinetown KL, Sherwood N, Gong S, Tetu SG, Midgley DJ, Paulsen IT. 2019. Succession Patterns and Physical Niche Partitioning in Microbial Communities from Subsurface Coal Seams. iScience 12:152–167.

53. Rosewarne CP, Greenfield P, Li D, Tran-Dinh N, Bradbury MI, Midgley DJ, Hendry P. 2013. Draft Genome Sequence of *Clostridium* sp. Maddingley, Isolated from Coal-Seam Gas Formation Water. Genome Announc 1:e00081–12.

54. Schink B. 2006. Syntrophic Associations in Methanogenic Degradation, p. 1–19. In Overmann, J (ed.), Molecular Basis of Symbiosis. Springer, Berlin, Heidelberg.

55. Rezaei Somee M, Dastgheib SMM, Shavandi M, Ghanbari Maman L, Kavousi K, Amoozegar MA, Mehrshad M. 2021. Distinct microbial community along the chronic oil pollution continuum of the Persian Gulf converge with oil spill accidents. 1. Sci Rep 11:11316.

56. Vick SHW, Greenfield P, Willows RD, Tetu SG, Midgley DJ, Paulsen IT. 2019. Subsurface Stappia: Success Through Defence, Specialisation and Putative Pressure-Dependent Carbon Fixation. Microb Ecol 80:34–46.

57. Zengler K, Richnow HH, Rosselló-Mora R, Michaelis W, Widdel F. 1999. Methane formation from long-chain alkanes by anaerobic microorganisms. 6750. Nature 401:266–269.

58. Grime JP. 1977. Evidence for the existence of three primary strategies in plants and its relevance to ecological evolutionary theory. Am Nat 111:1169–1194.

59. Bik EM, Costello EK, Switzer AD, Callahan BJ, Holmes SP, Wells RS, Carlin KP, Jensen ED, Venn-Watson S, Relman DA. 2016. Marine mammals harbor unique microbiotas shaped by and yet distinct from the sea. 1. Nat Commun 7:10516.

60. Storey MA, Andreassend SK, Bracegirdle J, Brown A, Keyzers RA, Ackerley DF, Northcote PT, Owen JG. 2020. Metagenomic Exploration of the Marine Sponge Mycale hentscheli Uncovers Multiple Polyketide-Producing Bacterial Symbionts. mBio 11:e02997–19.

61. Robbins SJ, Singleton CM, Chan CX, Messer LF, Geers AU, Ying H, Baker A, Bell SC, Morrow KM, Ragan MA, Miller DJ, Forêt S, Voolstra CR, Tyson GW, Bourne DG. 2019. A genomic view of the reef-building coral Porites lutea and its microbial symbionts. 12. Nat Microbiol 4:2090–2100.

62. Wang W, Tao J, Yu K, He C, Wang J, Li P, Chen H, Xu B, Shi Q, Zhang C. 2021. Vertical Stratification of Dissolved Organic Matter Linked to Distinct Microbial Communities in Subtropic Estuarine Sediments. Front Microbiol 12:697860.

